# Aging reshapes odor-evoked dynamics in central olfactory circuits

**DOI:** 10.64898/2026.09.30.755620

**Authors:** Ian F. Chapman, Kevin A. Bolding

## Abstract

Olfactory function declines with age, and olfactory impairment predicts subsequent cognitive decline and neurodegenerative disease. Age-related impairments in odor identification and discrimination implicate central olfactory circuits in both mice and humans, but it remains unclear where in the processing hierarchy aging alters neural function. To resolve this, we acquired simultaneous Neuropixels recordings from olfactory bulb (OB) and piriform cortex (PCx) in young and old mice during awake, head-fixed odor presentation, allowing us to examine successive processing stages within individual animals. Our recordings revealed a shift toward greater excitation in both regions and reduced inhibitory neuron activity in PCx. Aging also restructured odor-evoked local field potential (LFP) oscillations within and across regions, with strongly elevated beta power and coherence. Population odor representations were less discriminable in OB but not PCx, constituting the clearest neural analog of previously reported discrimination deficits. Single-unit, LFP, and population measures did not collapse onto a single axis using dimensionality reduction, indicating that several partly independent processes reshape central olfactory dynamics with aging. Together, these findings identify central circuit excitation/inhibition and oscillatory imbalances that parallel age-related olfactory dysfunction despite largely preserved odor responsivity.

## Introduction

Olfactory dysfunction is among the earliest markers of aging-related cognitive decline and neurodegenerative disease in humans^1–4^. This dysfunction is not simply a loss of sensitivity, but instead commonly presents as deficits in higher-order olfactory functions, such as identification and memory^3,5^. These higher-order functions are thought to depend on neural population representations within the olfactory bulb (OB) and downstream olfactory cortical areas, including the piriform cortex (PCx), the largest recipient of OB projections^6–11^. However, few studies have examined functional effects of normal aging on these central olfactory circuits. Rodent models have begun to address this gap, as odor-guided behavior is similarly impaired in aged mice, with the strongest and earliest effects on performance found in odor discrimination tasks^12–14^. While olfactory sensory neuron (OSN) responses are relatively preserved with aging in mice^15^, LFP recordings indicate altered rhythmic organization in the OB^16^, and recordings from *ex vivo* preparations indicate greater excitability in PCx neurons and reduced NMDAR-dependent plasticity in PCx of aged rats^17,18^.

Age-related changes in neural circuit function that might impact cognition and perception are heterogeneous, depending on species, brain region, and modality. However, some physiological changes, such as elevated firing rates during ongoing and stimulus-evoked activity, are repeatedly observed across multiple hippocampal and sensory cortical networks^19–22^. These effects may arise due to altered intrinsic excitability^23^ or weakened inhibitory regulation via loss of GABAergic cells, synapses, or inhibitory neural activity^24–28^. Moreover, neurons in aged somatosensory, visual, and auditory areas have reduced stimulus selectivity^19,29,30^, while place cell representations in the CA3 region of the aged hippocampus are less context-selective and carry less spatial information^20,31^. Notably, interventions intended to restore inhibitory tone improved stimulus selectivity in macaque V1^32^, and interventions targeting excess CA3 activity improve memory in aged rats^33^. Thus, age-dependent changes in the balance of excitation and inhibition are linked to altered neural coding and may, in turn, affect cognitive or perceptual function.

Whether or not this is the case in central olfactory structures remains unclear. Impaired excitatory and inhibitory interactions could underlie changes in oscillatory networks^16^, but this possibility has not been explored to this point *in vivo*. Furthermore, the impact of aging on central olfactory stimulus representations, a core function of the OB and PCx, has not been examined. To define how normal aging alters central olfactory circuit physiology and computation *in vivo*, we therefore recorded large populations of individual OB and PCx neurons simultaneously in young and aged mice during head-fixed, awake odor presentation. Our data reveal multiple indicators of altered circuit dynamics across central olfactory structures, including increased odor-evoked activity, divergent local field potential dynamics, and impaired stimulus separability in the OB. Together, these features indicate that normal aging produces broad, multivariate changes in odor-evoked dynamics of central olfactory circuits.

## Results

### Similar odor responsivity in the OB and PCx of young and old mice

To examine effects of age on central olfactory processing, we acquired simultaneous recordings using Neuropixels 2.0 probes positioned in OB and PCx (Fig. 1a) of young (n = 7, 3–4 months) and old (n = 9, 19–26 months) mice in an awake, head-fixed preparation during presentation of a panel of 15 monomolecular odorants at 0.5% v/v dilution (Supplementary Table 1). While likely higher than values found in natural settings^34^, this concentration was selected to be well above detection threshold for mice^35^ to ensure epithelial activation. This age range in mice approximately corresponds to ages 50–75 in humans^36^, and precedes significant increases in mortality and prevalence of systemic health issues in mice. Spike sorting yielded large populations of stable, well-isolated units (Supplementary Table 2, OB young: n = 534, mean = 76.3 ± 12.3 SEM units per session; OB old: n = 624, 69.3 ± 10.4; PCx young: n = 2551, 364.4 ± 46.5; PCx old: n = 2402, 300.3 ± 44.6) spanning dorsal and ventral OB and all layers of PCx with spatial distributions consistent with elevated densities in the OB mitral cell layer and PCx layer 2 (Supplementary Fig. 1). Respiration was monitored with a flow sensor positioned across from the nose. Odor delivery was triggered on exhalation and odor responses were aligned to the start of the following inhalation for analysis (Fig. 1b). Consistent with previous reports^8,10,11^, individual OB and PCx cells in both age groups could produce temporally precise spiking responses following odor inhalation with heterogeneous tuning selectivity and response durations (Fig. 1b).

**Figure 1.**
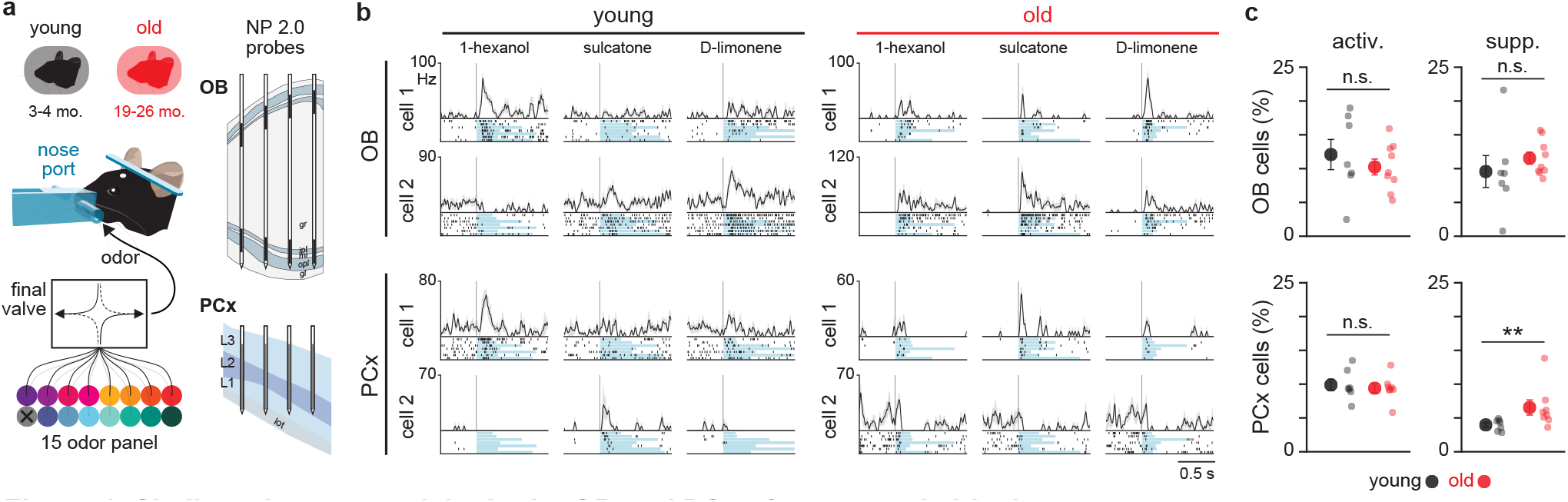
Similar odor responsivity in the OB and PCx of young and old mice. **a** Schematic of the recording setup. Young (3–4 months) and old (19–26 months) mice are head-fixed with a custom nose port to monitor respiration and deliver odor stimuli from a multichannel olfactometer. Four-shank Neuropixels 2.0 probes are inserted acutely into the OB and the PCx on the day of recording. **b** Example odor response rasters and PSTHs from two individual OB and PCx cells in young and old mice, showing responses to three odorants. Blue shading indicates odor presentation. **c** Percentage of cells activated or suppressed by odor stimuli in OB or PCx of young (black) and old (red) animals. Large circles and error bars indicate mean ± SEM; small points show individual animals. Values show mean ± SEM across animals. **p < 0.01; two-sided Wilcoxon rank-sum test.

Odor sensitivity declines with age in mice^13^ and humans^37,38^. We therefore asked whether central olfactory neural populations are less responsive to odors in old mice. The percentage of cells significantly activated by odors did not differ between young and old mice in either OB or PCx (Fig. 1c, OB: n = 7 young and 9 old mice, p = 0.368; PCx: n = 7 young and 8 old mice, p = 0.908; rank-sum tests). The percentage of cells suppressed by odor was also indistinguishable in young and old OB populations (Fig. 1c, n = 7 young and 9 old mice, p = 0.101; rank-sum test). However, there was a significant increase in the percentage suppressed by odor in the old PCx (Fig. 1c, n = 7 young and 8 old mice, p = 0.008; rank-sum test). Breathing rate during the first 2 s of odor delivery was not significantly different between young and old and was likely not a source of variation in responsivity (Supplementary Fig. 2). Overall, these results are not consistent with a general age-related loss of central responsivity to odors at the stimulus concentrations used in our experiments.

### Hyperactivity in the OB and PCx of aged mice

Cortical networks exhibit increased excitability and spiking activity during aging^19,20,22,23^, but age-related activity changes have not been described in the olfactory system *in vivo*. To address this, we examined activity in large populations of simultaneously recorded OB neurons during odor presentation (Fig. 2a). We first considered overall firing rates for each recorded animal (Fig. 2b). We found that average firing rates were not significantly different between young and old (Fig. 2b, n = 7 young and 9 old mice, p = 0.315, rank-sum test). There was also no difference in the distribution of overall firing rates across all cells recorded from the young and old OB (Fig. 2b, n = 534 young and 624 old cells, p = 0.358, Kolmogorov-Smirnov test). To examine OB activity during odor delivery trials, we averaged population firing rates during a pre-odor baseline breath and in the first breath after odor delivery. Activity during baseline breaths was significantly elevated in the old OB (Fig. 2c, n = 105 young and 135 old population-odor pairs, p < 0.001, rank-sum test), but odor-evoked activity did not differ between young and old (n = 105 young and 135 old population-odor pairs, p = 0.145, rank-sum test). However, responses visualized as population-averaged peristimulus time histograms (PSTHs) clearly differed between age groups in the strength of their initial odor-evoked rise (Fig. 2d). The peak response in this early period was reliably larger for old OB populations compared to young (Fig. 2d, n = 105 young and 135 old population-odor pairs, p < 0.001, rank-sum test). Thus, aging effects on OB activity emerged after respiration alignment, with higher baseline-breath activity and a substantially stronger odor-evoked population peak.

**Figure 2.**
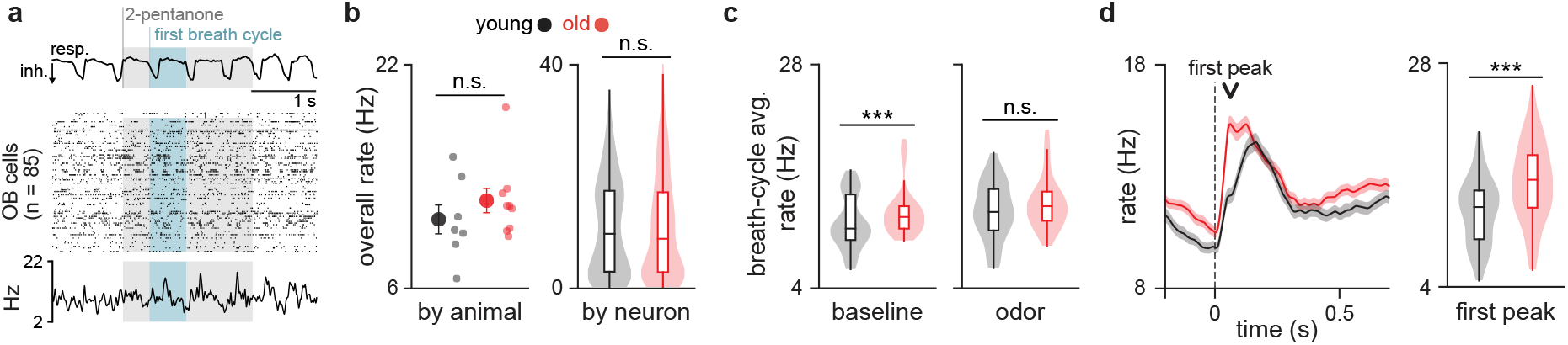
Hyperactivity in the OB of old mice. **a** Example respiration trace, population raster, and population-averaged firing rate from a single odor trial in the OB of a young animal. Blue shading indicates the first respiration cycle after odor delivery. **b** Whole-session firing rates of OB cells, averaged by animal (left) or shown across individual cells (right). **c** Average OB firing rate for each population-odor pair during a pre-odor baseline respiration cycle or the first respiration cycle after odor delivery. **d** Inhalation-aligned population PSTHs during odor presentation. Downward arrow indicates the time used to quantify the initial response peak. PSTHs show mean ± SEM across population-odor pairs; violin plots show the distribution of peak responses across population-odor pairs. For measures averaged by animal, mean is shown with SEM as error bars. Strip plots to the right of each mean show individual datapoints. Violin plots show the distribution of values, with embedded box plots indicating the median and interquartile range. Animal-level and population-odor pair comparisons were performed using two-sided Wilcoxon rank-sum tests; comparisons of firing-rate distributions across individual cells in **b** used two-sample Kolmogorov–Smirnov tests. ***p < 0.001.

We next asked whether this increased excitation in the OB was mirrored in downstream PCx activity. Given evidence of decreased inhibitory tone with aging in other cortical systems^23–26,39,40^, we further characterized our data by identifying putative inhibitory neurons in our extracellular recordings. We first clustered units using our larger database of Neuropixels 2.0-recorded PCx units (n = 61 recordings, n = 19,791 total units) based on waveform and spiking features (Supplementary Fig. 3). To anchor this analysis, we used putative inhibitory interneurons identified by opto-tagging in separate PCx recordings from PV-Cre::Ai32 animals (Fig. 3a,b; n = 9 animals, n = 37 total cells) as well as cross-correlogram analysis to identify putative outgoing excitatory (EXC_CCG_) and inhibitory (INH_CCG_) monosynaptic interactions. This method consistently produced a cluster enriched for opto-tagged PV cells over a wide range of embedding and clustering parameters (Supplementary Fig. 4). We therefore selected final clustering parameters that minimized EXC_CCG_ units in the PV-enriched cluster while maximizing overall clustering quality via silhouette score. The final PV-enriched cluster contained the majority of INH_CCG_ units (Fig. 3c) and comprised 7.85 ± 0.77% of cells in our young and old PCx recordings (n = 15 recordings). Units assigned to this cluster are referred to as putative PCx inhibitory cells (PC_INH_), whereas all remaining units are operationally referred to as putative PCx excitatory cells (PC_EXC_).

**Figure 3.**
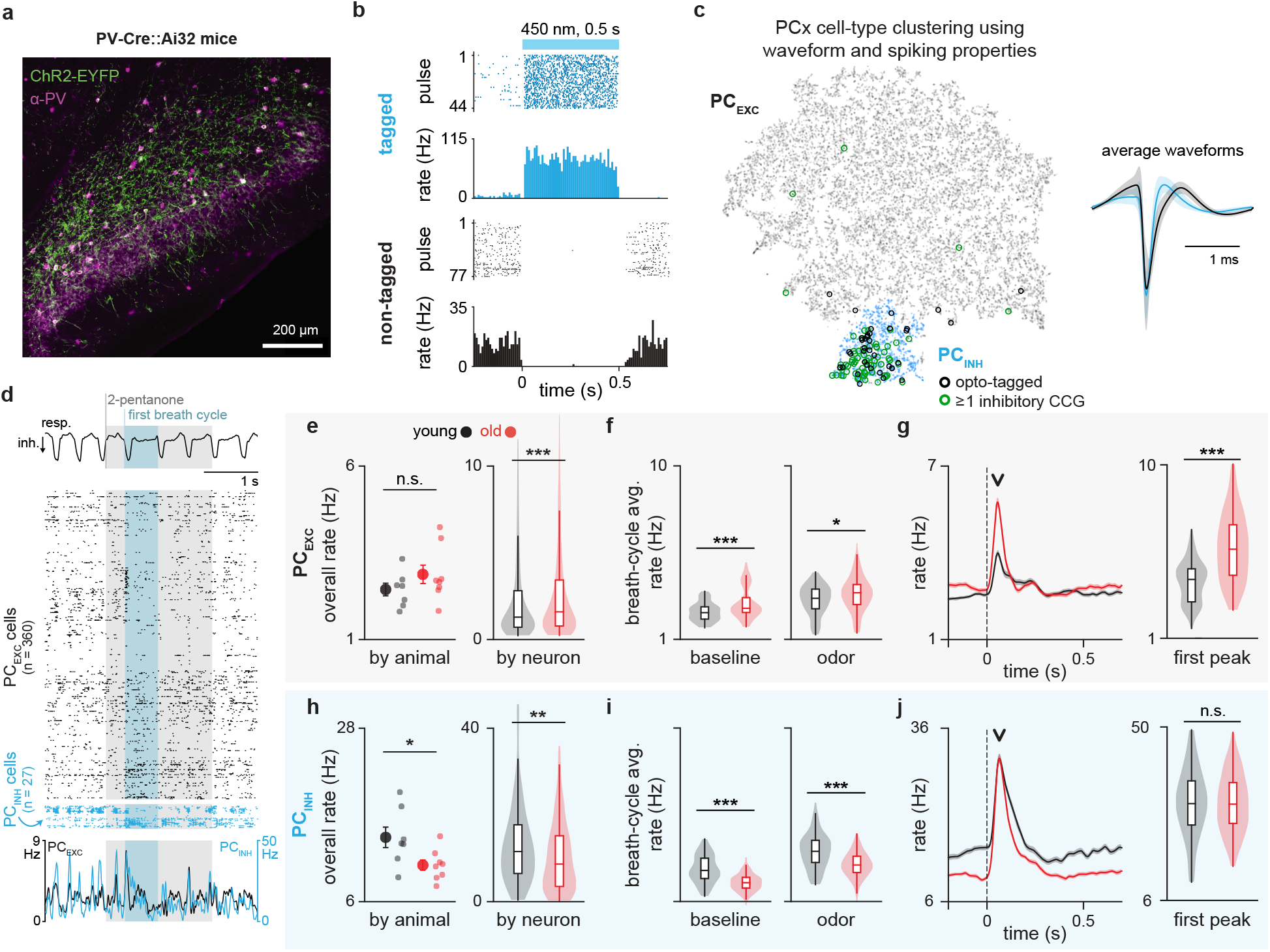
Altered excitatory and inhibitory activity in the PCx of old mice. **a** Representative PCx section from a PV-Cre::Ai32 animal showing ChR2-EYFP and PV immunolabeling used to validate optogenetic targeting. **b** Example raster and PSTH plots for a tagged and non-tagged unit during light stimulation. **c** Left, t-SNE projection of PCx units based on waveform and spiking properties, with colors indicating DBSCAN cluster assignments. Units assigned to the PC_INH_ cluster are shown in blue and all remaining units are designated PC_EXC_. Opto-tagged units and units with putative inhibitory CCG interactions are indicated by overlaid markers. Right, average waveforms of PC_EXC_ and PC_INH_ cells. **d** Example respiration trace, population rasters, and population-averaged firing rates from simultaneously recorded PC_EXC_ and PC_INH_ populations during a single odor trial in a young animal. Blue shading indicates the first respiration cycle after odor delivery. **e, h** Whole-session firing rates of PC_EXC_ (**e**) and PC_INH_ (**h**) cells, averaged by animal (left) or shown across individual cells (right). **f, i** Average firing rates for each population-odor pair during a pre-odor baseline respiration cycle or the first respiration cycle after odor delivery for PC_EXC_ (**f**) and PC_INH_ (**i**) cells. **g, j** Inhalation-aligned population PSTHs and initial peak responses for PC_EXC_ (**g**) and PC_INH_ (**j**) cells. Downward arrows indicate the times used to quantify the initial response peaks. PSTHs show mean ± SEM across population-odor pairs; violin plots show the distribution of peak responses across population-odor pairs. For measures averaged by animal, mean is shown with SEM as error bars. Strip plots to the right of each mean show individual datapoints. Time-resolved plots show mean ± SEM. Violin plots show the distribution of values, with embedded box plots indicating the median and interquartile range. Animal-level and population-odor pair comparisons were performed using two-sided Wilcoxon rank-sum tests; comparisons of firing-rate distributions across individual cells in **e** and **h** used two-sample Kolmogorov–Smirnov tests. *p < 0.05, **p < 0.01, ***p < 0.001.

We next examined activity in simultaneously recorded populations of putative PCx excitatory and inhibitory neurons during odor presentation (Fig. 3d). Average whole-session firing rates did not differ significantly between young and old PC_EXC_ cells at the animal level (Fig. 3e, n = 7 young, 8 old; p = 0.298, rank-sum test), but were significantly lower in old PC_INH_ cells (Fig. 3h, n = 7 young, 8 old; p = 0.049, rank-sum test). Across all recorded cells, firing-rate distributions differed significantly between age groups for both PC_EXC_ cells (Fig. 3e, n = 2370 young, 2182 old; p < 0.001, Kolmogorov-Smirnov test) and PC_INH_ cells (Fig. 3h, n = 181 young, 220 old; p = 0.003, Kolmogorov-Smirnov test). These distributional changes had opposite directions, with PC_EXC_ firing rates shifted toward higher values in old mice and PC_INH_ firing rates shifted toward lower values. Firing rates averaged over single breaths during baseline and in the first respiration cycle after odor delivery were significantly increased in PC_EXC_ (Fig. 3f, n = 105 young and 120 old population-odor pairs, baseline: p < 0.001; odor: p = 0.014, rank-sum tests), and significantly decreased in PC_INH_ populations (Fig. 3i, n = 105 young and 120 old population-odor pairs, baseline: p < 0.001; odor: p < 0.001, rank-sum tests). As in the OB, responses in PC_EXC_ cells differed most strongly between age groups during the early response peak after odor inhalation (Fig. 3g, n = 105 young and 120 old population-odor pairs, p < 0.001, rank-sum test). By contrast, the peak of odor-evoked responses in PC_INH_ cells was not significantly different between age groups (Fig. 3j, n = 105 young and 120 old population-odor pairs, p = 0.772, rank-sum test). Thus, although PC_INH_ activity was lower in aged PCx during baseline and breath-averaged odor responses, the output of the inhibitory population reached similar peak firing rates early in the odor response. To determine whether reduced inhibitory activity was accompanied by a loss of inhibitory neurons, we quantified the density of parvalbumin-positive (PV+) cells in anterior PCx from the same aging cohort. PV+ cell density did not differ significantly between young and old mice, either across the full cortical depth or within individual cortical layers (Supplementary Fig. 5). Taken together, these results indicate a shift towards increased excitatory and decreased inhibitory activity with age in central olfactory circuits, without a detectable reduction in PV+ neuron density.

### Increased odor-evoked beta power and coherence in the OB and PCx of aged mice

Balanced excitatory and inhibitory interactions govern coordinated network activity in sensory circuits, expressed as local field potential (LFP) oscillations. To ask whether the spike-level changes observed in aged animals were accompanied by altered network dynamics, we examined odor-evoked LFP spectra in OB and PCx. Odor inhalation evoked prominent beta (10–30 Hz) and gamma (30–100 Hz) frequency oscillations in both OB and PCx as previously described^41–43^ (Fig. 4a). Marked age-dependent differences in both beta and gamma power were apparent in both OB and PCx (Fig. 4a–c). Inspection of odor-evoked spectrograms revealed distinct patterns of activity above and below 50 Hz within the gamma range (Fig. 4b). We therefore divided gamma into low (30–50 Hz) and high (50–100 Hz) frequency bands for subsequent analyses. Baseline beta power did not differ between groups in the OB or the PCx (Fig. 4d, OB n = 105 young and 135 old population-odor pairs, p = 0.169, rank-sum test; PCx n = 105 young and 120 old, p = 0.298, rank-sum test). However, odors evoked a large and sustained increase in beta power in old animals from both structures (Fig. 4d, OB n = 105 young and 135 old population-odor pairs, p < 0.001, rank-sum test; PCx n = 105 young and 120 old, p = 0.01, rank-sum test) particularly after the first 100 ms (Fig. 4c).

**Figure 4.**
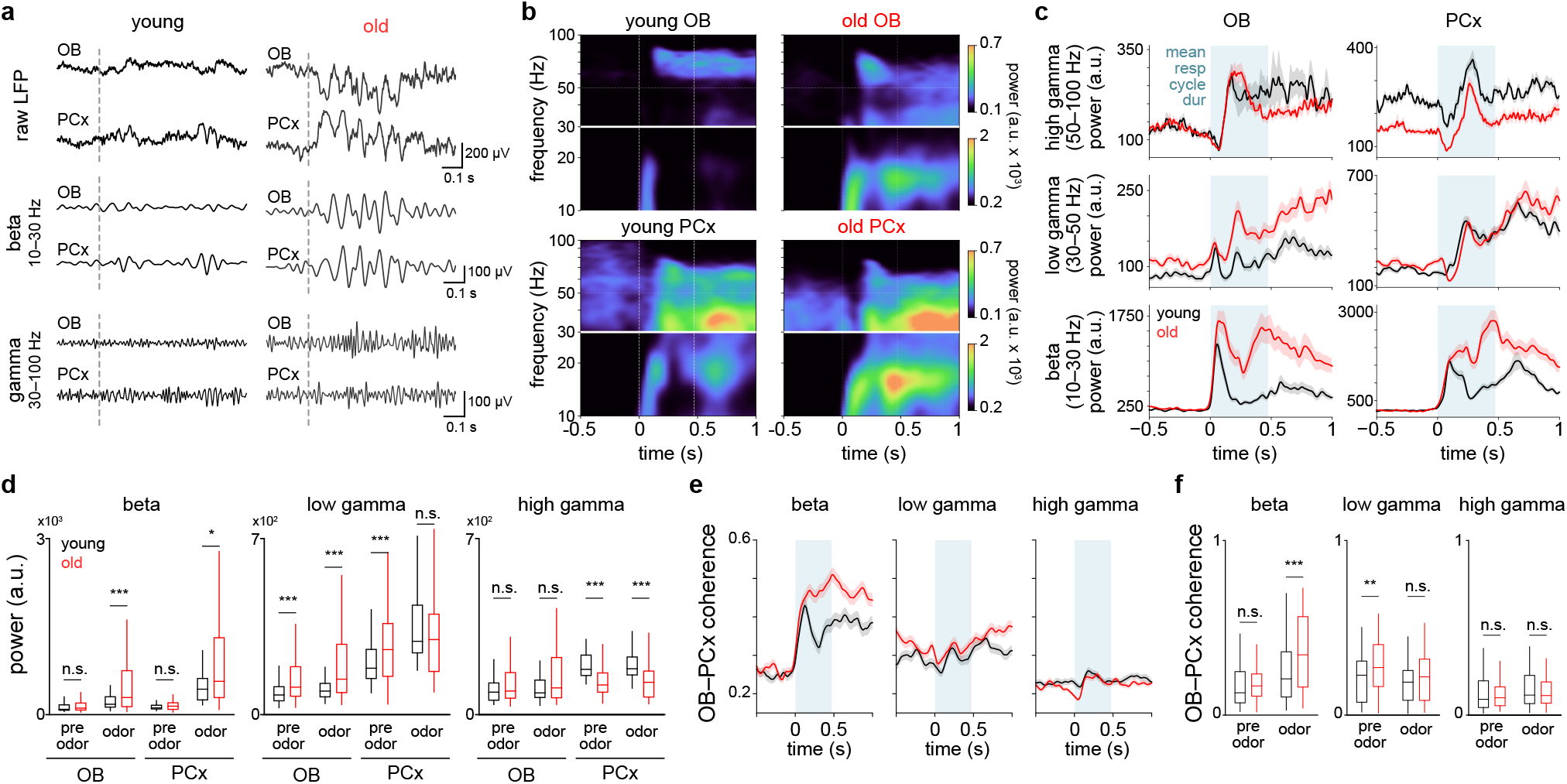
Increased odor-evoked beta power and coherence in OB and PCx of old mice. **a** Example LFP traces from simultaneous OB and PCx recordings in a young and old animal during a single odor trial. Traces show raw LFP or signals bandpass filtered in the beta (10–30 Hz) or gamma (30–100 Hz) ranges. **b** Inhalation-aligned spectrograms averaged across population-odor pairs for young and old OB and PCx. Spectrogram power was whitened by multiplying by frequency for visualization. Separate color scales are used for the beta (10–30 Hz) and gamma (30–100 Hz) frequency ranges. **c** Time-resolved beta (10–30 Hz), low-gamma (30–50 Hz), and high-gamma (50–100 Hz) power during odor stimulation, averaged across population-odor pairs in OB and PCx. Blue shading indicates the 475-ms analysis window, corresponding to the mean first respiration-cycle duration. **d** Beta, low-gamma, and high-gamma power in OB and PCx during 475-ms baseline and odor-response windows. **e** Time-resolved OB–PCx coherence in the beta, low-gamma, and high-gamma bands, averaged across population-odor pairs. **f** OB–PCx coherence in the three frequency bands during the same 475-ms baseline and odor-response windows. Time-resolved plots show mean ± SEM. Violin plots show the distribution of values, with embedded box plots indicating the median and interquartile range. Population-odor pair comparisons were performed using two-sided Wilcoxon rank-sum tests. *p < 0.05, **p < 0.01, ***p < 0.001.

Low-gamma activity exhibited broader temporal dynamics that differed between age groups, whereas high-gamma power peaked approximately 200–300 ms after inhalation in both OB and PCx (Fig. 4c). Baseline and odor-evoked low-gamma power were significantly increased in the old OB (Fig. 4d, baseline n = 105 young and 135 old population-odor pairs, p < 0.001; odor-evoked p < 0.001, rank-sum tests). Neither baseline nor odor-evoked high-gamma power differed significantly between age groups in the OB (Fig. 4d, baseline p = 0.218; odor-evoked p = 0.184, rank-sum tests). In PCx, baseline low-gamma power was significantly increased in old mice, but odor-evoked low-gamma power did not differ significantly between age groups (Fig. 4d, n = 105 young and 120 old population-odor pairs, baseline p < 0.001; odor-evoked p = 0.122, rank-sum tests). By contrast, both baseline and odor-evoked high-gamma power were significantly lower in PCx of old mice (Fig. 4d, baseline p < 0.001; odor-evoked p < 0.001, rank-sum tests).

Because cross-regional communication between OB and PCx is thought to play a critical role in odor processing^42,44–46^, we next leveraged the simultaneously recorded signals in both regions to compute coherence between OB and PCx within the beta, low-gamma, and high-gamma bands (Fig. 4e). Beta coherence was significantly increased during odor responses in old mice (Fig. 4f, baseline n = 105 young and 120 old population-odor pairs, p = 0.095; odor-evoked p < 0.001, rank-sum tests). Baseline low-gamma coherence was significantly increased in old mice (Fig. 4f, n = 105 young and 120 old population-odor pairs, p = 0.002), but odor-evoked coherence did not differ significantly between age groups (p = 0.053, rank-sum tests). High-gamma coherence did not differ between age groups during baseline or odor responses (Fig. 4f, baseline p = 0.985; odor-evoked p = 0.563, rank-sum tests). Together, these results indicate that aging alters odor-evoked network dynamics in a band- and region-dependent manner, including stronger beta-band modulation within OB and PCx, increased beta coherence between these structures, and reduced high-gamma power in PCx.

### Impaired odor stimulus separability in the OB, but not PCx, of aged mice

Neural populations in OB and PCx represent odor identity using reliably distinguishable population codes thought to support odor discrimination and identification, olfactory abilities that decline with age. We therefore asked whether the discriminability of stimulus-specific population responses in the first sniff cycle after odor delivery was impacted by aging in OB or PCx. For PCx population analyses, we excluded putative inhibitory neurons (PC_INH_) and defined population response vectors using only excitatory projection neurons (PC_EXC_), the principal output neurons of the PCx network. We first visualized response similarity using Pearson’s correlation coefficients between trial-averaged odor response vectors from OB or PC_EXC_ populations pooled across recordings (Fig. 5a). Responses to different odors appeared markedly more similar in old OB than in young OB. To quantify stimulus separability, we compared trial-to-trial response correlations within and across odors within each session. The difference between within-odor and across-odor correlations (Δ correlation) was significantly lower in old OB (Fig. 5b, n = 105 young and 135 old, p < 0.001, rank-sum test), whereas no difference was observed in PCx (Fig. 5b, n = 105 young and 120 old, p = 0.505, rank-sum test).

**Figure 5.**
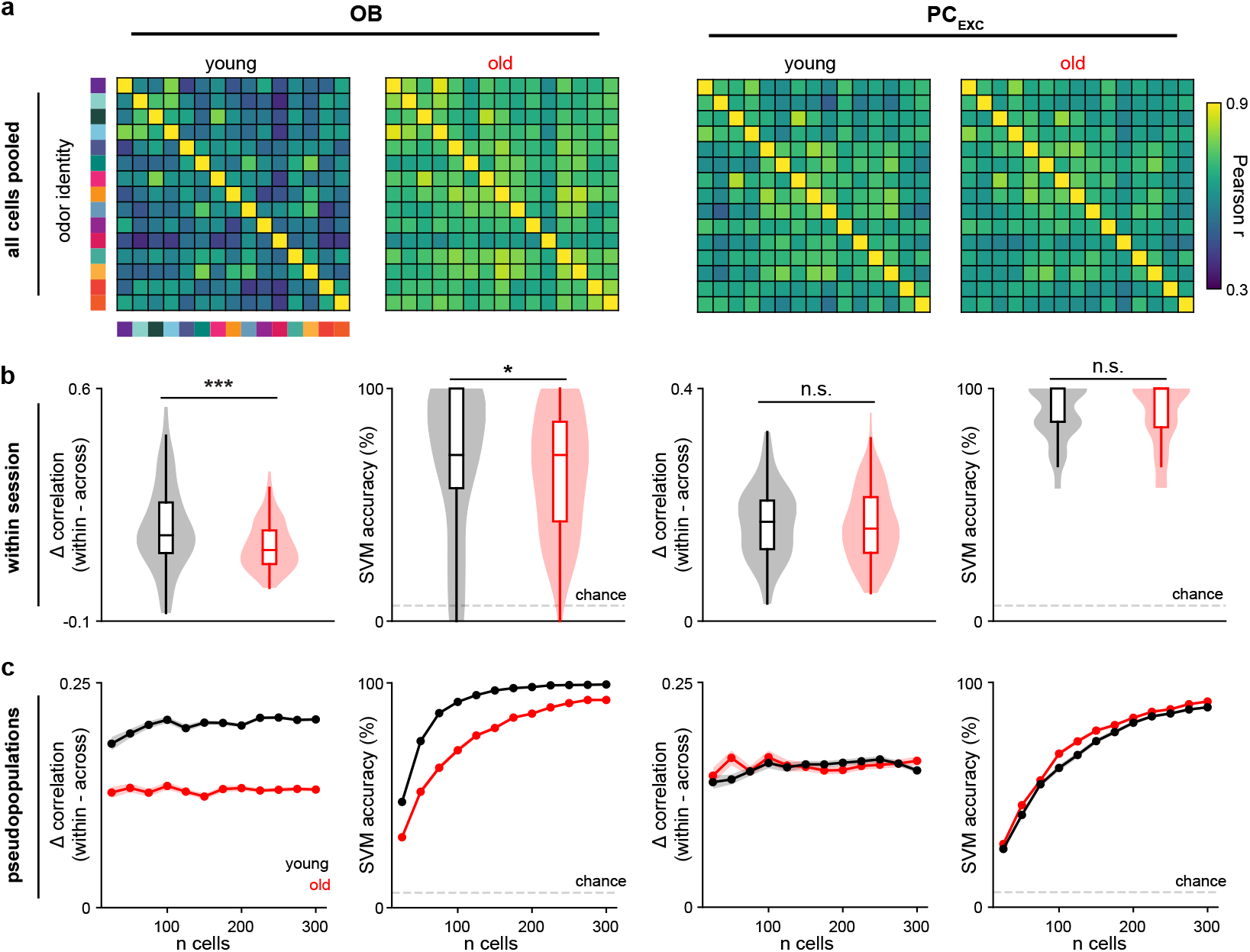
Impaired odor stimulus separability in the OB, but not PCx, of old mice. **a** Pearson correlation coefficients between trial-averaged odor response vectors from pooled OB or PC_EXC_ populations in young and old mice. **b** Δ correlation (within odor correlation − across odor correlation) and cross-validated SVM decoding accuracy for population-odor pairs within individual recording sessions. Violin plots show the distribution of population-odor pair values. Embedded box plots show the median and interquartile range. **c** Δ correlation and cross-validated SVM decoding accuracy as in **b**, but using random pseudopopulations of cells pooled across recordings. Fifty iterations were performed at each population size; lines show the mean across iterations with SEM as shading. Population-odor pair comparisons in **b** were performed using Wilcoxon rank-sum tests. *p < 0.05, **p < 0.01, ***p < 0.001.

To ask whether odor identity codes could be reliably distinguished by a downstream observer, we tested the accuracy of linear support vector machine (SVM) classifiers trained with first-sniff odor population responses. Cross-validated decoding accuracy was significantly lower using population responses from the old OB (Fig. 5b, n = 105 young and 135 old, p = 0.033, rank-sum test). In PCx, decoding accuracy remained similarly high in young and old mice (n = 105 young and 120 old, p = 0.754, rank-sum test). Total recorded cell yields did not differ between age groups (OB n = 7 young and 9 old, p = 0.673; PCx n = 7 young and 8 old, p = 0.337, Welch’s t-test) suggesting that differences in cell number were unlikely to explain the reduced OB decoding accuracy. To further assess how stimulus separability scaled with population size, we calculated Δ correlation and SVM accuracy using randomly selected pseudopopulations of OB and PCx cells across a fixed range of cell numbers (Fig. 5c). Decoding accuracy was lower in the old OB but not old PCx (Fig. 5c). Together, these analyses indicate that aging impairs odor stimulus separability in the OB, while odor identity representations in PCx remain comparatively preserved.

### Broader tuning of individual cells in OB and PCx of aged mice

To clarify what features of the neural population response contribute to the decrease in separability found in the aged OB, we first examined first breath response tuning using lifetime sparseness (Fig. 6a). Sparseness values near 1 indicate responses concentrated in a single stimulus, whereas values near 0 indicate similar responses across stimuli; lower values with age would therefore indicate broader tuning across odors. Per-animal averaged lifetime sparseness did not differ between young and old OB (n = 7 young and 9 old mice, p = 0.427, rank-sum test) or PCx (n = 7 young and 8 old mice, p = 0.418, rank-sum test). However, the distribution of lifetime sparseness values across cells differed significantly between age groups in both OB (n = 534 young and 624 old cells, D = 0.106, p = 0.003, Kolmogorov-Smirnov test) and PCx (n = 2370 young and 2182 old cells, D = 0.042, p = 0.034, Kolmogorov-Smirnov test; Fig. 6a), with old mice showing a shift toward lower lifetime sparseness values. In addition to changes in tuning breadth, larger odor-evoked responses in aged animals could arise from broader recruitment across the recorded population, increasing overlap between odor representations. We therefore calculated population sparseness for each population-odor pair as a measure of response density (Fig. 6b). Population sparseness did not differ between young and old animals in OB (n = 105 young and 135 old population-odor pairs, p = 0.084, rank-sum test) or PCx (n = 105 young and 120 old population-odor pairs, p = 0.288, rank-sum test), indicating similar response density between age groups.

**Figure 6.**
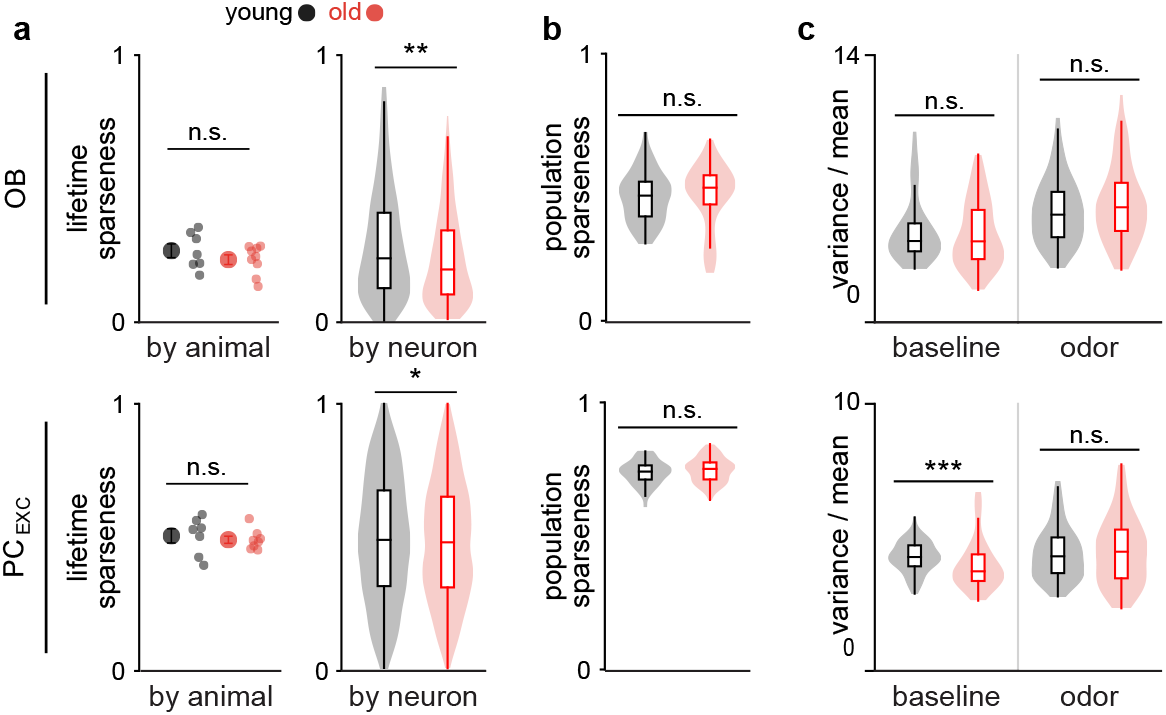
Broader tuning of individual neurons in OB and PCx of old mice. **a** Lifetime sparseness of OB and PC_EXC_ cells in young and old mice, shown as averages by animal (left) or distributions across individual cells (right). **b** Distribution of population sparseness of OB and PC_EXC_ odor responses across population-odor pairs. **c** Distribution of mean-adjusted variance of OB and PC_EXC_ firing rates during baseline and odor responses across population-odor pairs. Odor-response measures were calculated during the first respiration cycle after odor delivery. Violin plots show the distribution of values. Embedded box plots show the median and interquartile range. Animal-level and population-odor pair comparisons were performed using Wilcoxon rank-sum tests; comparisons of lifetime sparseness distributions across individual cells in a used two-sample Kolmogorov–Smirnov tests. *p < 0.05, **p < 0.01, ***p < 0.001.

Finally, since higher trial-to-trial variability might also explain decreased separability when using decoding approaches, we calculated variance during first breath responses, dividing by response mean to account for response magnitude (mean-adjusted variance, Fig. 6c). Baseline mean-adjusted variance (Fig. 6c) was significantly lower in the PCx (n = 105 young and 120 old population-odor pairs, p < 0.001, rank-sum test), but not in the OB (n = 105 young and 135 old population-odor pairs, p = 0.334, rank-sum test), indicating less variability in baseline PCx activity in the old animals. However, we did not find any significant differences in mean-adjusted variance during odor stimulation (Fig. 6c) in either the OB (n = 105 young and 135 old population-odor pairs, p = 0.172, rank-sum test) or the PCx (n = 105 young and 120 old population-odor pairs, p = 0.677, rank-sum test). Taken together, these results indicate that age-related changes in OB odor coding were most clearly associated with broader tuning in a subset of cells, while deficits in population separability occurred without global changes in response density or odor-evoked variability.

### Age-related changes are distributed across multiple dimensions of central olfactory physiology

Finally, we asked whether features of OB and PCx odor responses were predictive of age. We compiled 15 features that differed significantly with age in prior analyses and used a linear SVM to examine how these age-sensitive features jointly separated young and old animals. As expected from the feature-selection criterion, the combined feature set strongly separated age groups (93.3% leave-one-out classification accuracy; Fig. 7a). We therefore examined model weights to assess the relative contributions of individual features to this multivariate separation (Fig. 7b). The largest weights were assigned to E/I features (Fig. 2,3), with oscillatory measures (Fig. 4) and OB separability (Fig. 5) also contributing, consistent with the preceding analyses.

**Figure 7.**
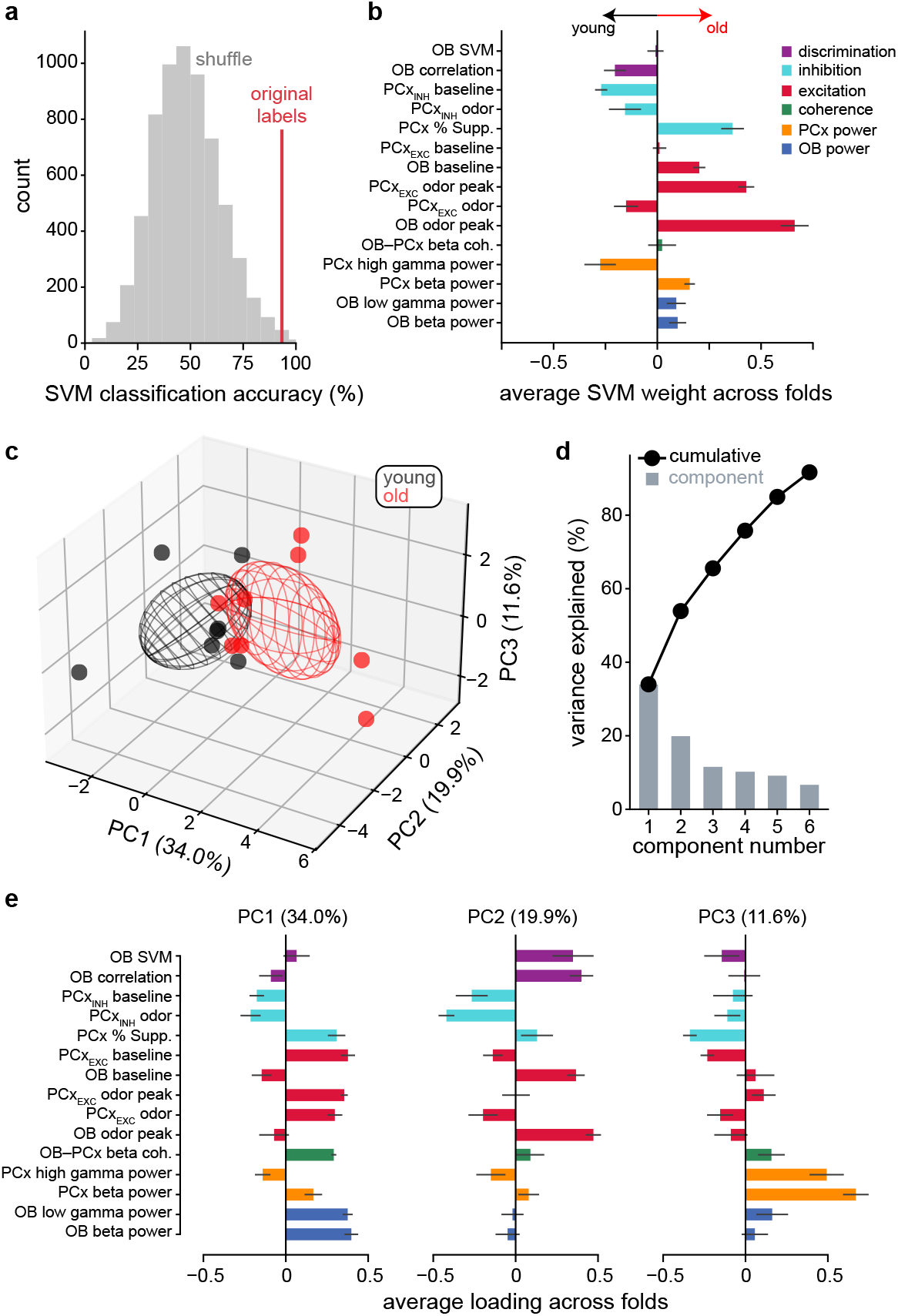
Age-related physiological changes are distributed across multiple dimensions of central olfactory circuit function. **a** Leave-one-out (LOO) SVM accuracy in predicting animal age (red line, 93.3%) compared with accuracies obtained from all possible permutations of age labels across animals (gray). **b** Feature weights from the SVM models in a, averaged across validation folds; error bars show SD. **c** Feature values projected onto the first three principal components. Points indicate individual animals and ellipsoids show the mean ± 1 SD for young (black) and old (red) animals. **d** Percent variance explained by each principal component after PCA on the curated feature set, with cumulative variance explained also shown. **e** Loadings for the first three principal components, averaged across LOO folds; error bars show SD.

We next used principal component analysis (PCA) to examine relationships between these features with age. Age group was apparent in the structure of the component projections (Fig. 7c), but the explained variance was distributed across multiple components (Fig. 7d), indicating that age did not map cleanly onto a single dimension of the data. Average component loadings across folds (Fig. 7e) showed that different age-sensitive features contributed to different dimensions. For example, measures of excitability (Fig. 2,3) and OB oscillatory power (Fig. 4) were strongly weighted on the first component, whereas OB separability (Fig. 5) contributed primarily to the second component and PCx oscillatory power (Fig. 4) to the third component. Thus, although central olfactory response features were sufficient to classify animal age with high accuracy, age-related physiological changes did not collapse onto a single shared axis. Instead, the data suggest that aging affects multiple partially independent dimensions of central olfactory physiology, consistent with contributions from more than one underlying aging process.

## Discussion

Aging leads to decline in higher-order olfactory abilities, such as identification and discrimination, that are thought to depend on population odor representations emerging in OB and downstream cortices. Our recordings demonstrate elevated excitation in both the OB and PCx, a reduction in PCx inhibition in old mice and reduced separability of odor representations in OB populations, with more subtle changes in tuning and sparseness in both structures. Overall odor-evoked activation was largely unaffected by age in both OB and PCx, consistent with previous reports that OSN input to OB is intact in aged mice . These results indicate that odor processing in central circuits is affected by age even when odor sensitivity is not grossly impaired.

In old animals, baseline activity was elevated in OB and PC_EXC_ cells, as were peak excitation and odor-evoked responses in PC_EXC_ cells. Conversely, PC_INH_ activity was reduced during ongoing activity and across much of the odor response, consistent with reduced inhibitory function in aging. These results agree with multiple studies showing decreases in inhibitory function with aging that increase excitability and impact function across a number of cortical structures^39,40,47–50^. However, the reduction in PC_INH_ neuron activity did not extend to the odor-evoked peak, which was similar in young and old mice, indicating that inhibitory neurons can still be strongly recruited by sensory input despite broader reductions in their activity. The preserved peak PC_INH_ response means that the larger odor-evoked excitatory peak in old mice cannot be explained simply by a corresponding reduction in peak inhibitory neuron recruitment. Moreover, we found no detectable reduction in PV+ neuron density in anterior PCx, suggesting that reduced inhibitory activity is not simply attributable to a loss of PV+ interneurons. One possibility is that reduced inhibitory restraint at odor onset permits a larger initial excitatory burst, which then recruits inhibitory neurons through local recurrent circuits, allowing inhibitory activity to reach a similar peak shortly thereafter. Alternatively, individual PCx excitatory cells may fire more strongly independent of inhibitory state due to age-related changes in intrinsic excitability^17^. With our current methods, we cannot distinguish these possibilities or fully resolve the underlying recurrent excitatory and inhibitory dynamics within PCx. Recordings from additional genetically and anatomically defined inhibitory cell types and temporally resolved computational modeling of E/I interactions within PCx may help identify the mechanisms of increased excitatory responses.

We find altered central oscillatory dynamics during odor processing with aging. Altered gamma oscillations have been previously reported in the OB with aging^16^, and increases in gamma power are likely driven by impaired GABA function in Alzheimer’s disease^51^. Gamma oscillations are generally associated with improved odor discrimination in the OB and PCx and are thought to reflect heightened local processing^43,52,53^. Thus, it is unexpected that increased low-gamma power in aged OB is paired with less distinguishable odor representations, and PCx odor identity coding is unaffected despite significantly weakened high-gamma oscillations. These results do not support a simple relationship between gamma oscillatory strength and the computational task of odor discrimination. Gamma oscillations in the aging olfactory system may reflect aberrant states that are not strongly related to their normal function in young OB and PCx. Alternatively, gamma oscillations may support computational tasks which were not assayed by our experiments and which could be revealed by recording in a more complex or demanding behavioral context.

Our recordings revealed a striking increase in beta oscillatory power and OB-PCx beta coherence with age. Central olfactory beta oscillations are strongly engaged during odor discrimination learning^41,42^ and depend on intact centrifugal feedback connectivity from PCx to OB^54^. Thus, increased beta power and coherence may indicate tighter functional coupling between OB and PCx in old animals. It is unclear whether this change is ultimately detrimental or beneficial for olfactory processing. Increased beta may reflect a compensatory change to address the reduced initial discriminability of OB odor representations. Alternatively, abnormally strong OB-PCx coupling and rhythmic patterning could come at the expense of local computations that depend on other frequency bands or asynchronous temporal organization.

Previous reports have repeatedly identified odor discrimination impairments in aged mice^12–14^, leading to the expectation that odor representations may be less separable in aged mice at the level of neural population coding. Here, we identify a candidate neural mechanism for reduced discrimination ability with aging; OB population responses are more correlated across odors and have a reduced ability to support accurate odor identity decoding. Surprisingly, this population coding deficit is not inherited by PCx. Thus, our results suggest that abnormalities in PCx excitation and temporal coordination are not sufficient to impact the separability of cortical odor representations during passive odor presentations. Previously described odor discrimination deficits were found under relatively challenging conditions, in which animals were required to make a fine discrimination or to learn multiple discrimination pairs. A limitation of our study is that our approach does not directly assay the animals’ discrimination ability for comparison with neural decoding outcomes. Additional challenges to the system during *in vivo* recordings, such as increasing demands on learning and memory, varying concentrations, or parsing complex mixtures may reveal consequences of altered PCx physiology that are not captured by simple odor identity decoding.

Prior studies across cortical systems during aging have linked reduced inhibition and increased excitability to reduced tuning selectivity and greater overlap in sensory representations^39,48^. We find that these features of cortical aging are only recapitulated piecemeal, rather than as a unified physiological phenotype, in the early central stages of olfactory processing. No one feature dominates predictive power for age group classification. Rather, SVM weights are distributed across features. Furthermore, our PCA results indicate that variance in age-related physiological variables is distributed across multiple lower-dimensional components instead of collapsing onto a single aging dimension encompassing concurrent changes in all physiological features. These observations indicate that normal aging does not have a unitary effect on olfactory processing but instead acts through multiple underlying processes. Thus, our results indicate that strategies for understanding age-related olfactory dysfunction will require mapping how distinct changes in excitability, stimulus coding, and temporal coordination within central olfactory networks contribute to behavioral and cognitive decline.

## Materials and Methods Mice

Experimental protocols were approved by the Monell Institutional Animal Care and Use Committee (IACUC). The primary aging dataset used male and female C57BL/6J (n = 16 total, Jackson Laboratory, # 000664) mice housed at Monell under standard conditions (12 hr. light/dark cycle, ad libitum chow and water). Animals were checked for basic mobility and health prior to surgical procedures and recordings. For opto-tagging experiments, male and female PV-Cre (+/+) mice (Jackson Laboratory, # 008069) were crossed with Ai32(+/+) mice (Jackson Laboratory, # 024109) to express ChR2 in parvalbumin positive cells (n = 9 mice total). Male and female C57BL/6J (n = 37) mice were used for additional PCx recordings added to the clustering dataset.

### Surgical procedures

Mice were anesthetized using isoflurane (5% induction and 2% maintenance) for all surgical procedures and given pre-operative analgesics subcutaneously (Meloxicam 5 mg/kg). The skin was removed over the dorsal skull, the skull was debrided, and the head was leveled using bregma and lambda midline landmarks. Recording sites were marked for the PCx (+1.32 mm anterior and 2.85 mm to the right of bregma) and ground pins were inserted into craniotomies above the cerebellum. A combination of first Metabond (Parkell Inc.), then cyanoacrylate glue (Bob Smith Industries), followed by conventional dental cement (Ortho-Jet) was used to fix the ground pins to the skull and to adhere a titanium bar for later head fixation. Animals were given analgesics (Meloxicam 5 mg/kg) for 3 days postoperatively and monitored for health and recovery. 24 hours prior to the first recording, mice were anesthetized using isoflurane as before and craniotomies were made for recording in both the OB and the PCx. PCx craniotomies were made beginning at the marked point and extending 1 mm posterior to allow for insertion of all 4 recording shanks. OB craniotomies were made over the dorsal surface of the right OB (ipsilateral to PCx craniotomies). Exposed dura was then covered with a silicone gel (DOWSIL™ 3-4680). Mice were monitored for any signs of pain or distress prior to recordings.

### Acute probe implantation and targeting

Animals were first acclimated to head-fixation and the nosecone across 3 days with sessions of increasing length (20, 40, and 60 minutes). On the day of recording, mice were head-fixed and probes painted with DiI or DiO for track labeling were gently inserted through the silicone gel above each craniotomy and slowly lowered into the brain at fixed speeds (<= 6 µm/s) using micromanipulators (Sensapex). For opto-tagging experiments, a custom probe with a tapered optic fiber (1 mm active length) attached parallel to the recording sites was used. PCx probes were inserted at a small (< 3°) outward angle from the sagittal plane to target PCx perpendicular to layer 2. OB probes were inserted ipsilateral to the PCx probes at a ∼20° inward angle relative to the sagittal plane to provide adequate spacing between the probes and manipulators during implantation. OB probes were initially inserted into the anterior and lateral quadrant of the dorsal surface to maximize spacing. During implantation, probes were monitored for both LFP signal and single unit spiking. In the OB, implantation was continued until dense spiking was observed from the ventral sites (∼2 mm depth from the surface depending on insertion site). When this was the case, dorsal sites were then scanned for spiking activity; if robust spikes were detected for at least 30 minutes, the channel map was reconfigured to maximize coverage of both dorsal and ventral OB. When dorsal spikes were not robust, sites were selected to maximize channel density in ventral regions. For the PCx, both LFP and spiking activity were closely monitored around the expected depth range (3.3-3.8 mm ventral from the surface). Large bands of dense spiking (> 50 µm in width) centered around a polarity reversal in LFP indicated adequate targeting to layer 2. PCx probes were lowered to maximize the number of shanks in PCx, but were stopped when any flutter or movement of the probes indicated a shank was nearing the ventral skull. Probes were then allowed to settle for at least 30 minutes prior to the start of a recording to minimize drift.

### Histological processing

After recordings, mice were deeply anesthetized with ketamine/xylazine (100/10 mg/kg) and transcardially perfused with 0.9% w/v saline followed by 10% neutral buffered formalin. Brains were post-fixed for 24 hours in formalin and cryoprotected in 30% sucrose before coronal sectioning. For validation of ChR2 expression in PV-Cre::Ai32 mice, 40 µm sections were stained with an anti-PV antibody (Synaptic Systems #195 308, 1:1000 dilution) and an anti-GFP antibody (Abcam #ab13970, 1:1000 dilution) to amplify EYFP-ChR2 expression. Secondary antibodies (Thermo Fisher #A-11076 and #A32931) were applied before acquiring merged images (Olympus BX63) to validate ChR2 expression in PV neurons.

For PV cell counting in anterior PCx of young and old mice, 12 µm sections were stained with the same anti-PV primary antibody and one of two secondary antibodies (Thermo Fisher #A-11076 or #A-11073). Images were acquired using the same microscopy system. PV-positive neurons were counted manually, and layer 1, 2, and 3 boundaries were delineated by an experimenter blinded to age group. Total PV+ cell counts and corresponding layer areas were summed across 1–6 sections per animal to calculate area-normalized PV+ cell density (cells/mm^2^). Per-animal densities were compared between age groups using two-sided Wilcoxon rank-sum tests.

### Data acquisition

Respiration data were monitored throughout the recording using an airflow sensor (Honeywell AWM3300V) mounted in the nosecone directly opposite the mouse’s nose. Electrophysiological data were acquired using 4 shank Neuropixels 2.0 probes and headstages from IMEC. Signals were sent to a dedicated acquisition card (IMEC PXIE_1000) in a NI PXIe-1083 chassis. A second card on the same chassis (PXIe-6341) was used to collect non-electrophysiological signals (respiration, odor triggers, opto-pulse times, etc.) for later alignment during analysis. Raw data were collected at 30 kHz in binary format using the OpenEphys GUI and Neuropixels PXI plugins, with any visualization used for probe monitoring/targeting performed after data were written to disk.

### Odor stimuli and delivery

15 monomolecular odor stimuli were chosen to span a variety of chemical functional groups (**Supplemental Table 1**). Odors were diluted to 0.5% concentration with caprylic/capric triglyceride oil (CCT, Spectrum Chemical) prior to loading onto the olfactometer. The olfactometer was based on prior designs in which 2 air paths (neutral/odor) were constantly maintained at 1 LPM flow rate using mass flow controllers (Alicat). Dedicated solenoids controlled when air was passed over a given odor vial and all odor vials were connected via a combination manifold (IDEX) with a downstream simultaneous final valve (NResearch) to control which air path was delivered to which downstream output (mouse/exhaust). On a given trial, odor was first equilibrated to bring the line to stable pressure/flow by turning on the solenoid for a given odor for 10 seconds. Then, after an exhalation was detected via respiration monitoring, the final valve was powered for 2 seconds to switch the air path delivered to the mouse to the odorized line prior to the next inhalation. In between trials, air on the odorized side of the olfactometer was passed through a blank (CCT only) vial to clean the olfactometer tubing. All tubing downstream of odors was made of PTFE to minimize odor adhesion in the lines between trials. Tubing around the nose was anchored using a nosecone milled from PTFE that directed odor perpendicular to the nose. A vacuum line opposite the odor entry side of the nosecone maintained flow across the nose and evacuated odors after delivery. Prior to delivery for experiments on any given recording day, each channel on the olfactometer was activated at least once during a calibration sequence to minimize differential pressure across olfactometer channels and confirm no leaks or malfunctioning hardware. Trial triggering and respiration monitoring were performed using custom code written in Bonsai Rx interfacing with dedicated Arduinos. TTL triggers for the final valve and the respiration data were also sent to the NI chassis to align with the electrophysiology data. During each recording, odors were delivered at fixed 30 s intervals in a block structure, where each stimulus was delivered once per block. This block structure was repeated with pseudorandomized trial order until 7 trials of each odor were delivered. Animals also received presentations of 2 or 4 component mixtures of the original 15 monomolecular stimuli in each block, but mixture data were not used for analysis in this manuscript.

### Local field potential analysis

Recordings were downsampled to 1 kHz for all LFP analysis, and signal processing was performed using the SciPy package in Python. For PCx probe localization, we applied a third-order Butterworth bandpass filter for gamma (30–100 Hz), then searched for periods of elevated gamma amplitude. Peaks in the gamma cycle during these periods were used to align and average broadband LFP segments (Supplementary Fig. 1). Each channel’s temporal mean was subtracted from the averaged segment, and the polarity-reversal depth was estimated by interpolating the voltage zero crossing across channels at the triggering peak. When no acceptable zero crossing was detected, RANSAC regression was used to estimate the reversal depth by extrapolation. For one animal, this approach found no valid shanks on the PCx probe, so only that animal’s OB probe was included for analysis.

For quantitative LFP analysis, PCx sites within 75 ± 25 µm ventral to the estimated polarity-reversal depth were averaged within each valid shank, followed by averaging across shanks to obtain one LFP trace per recording. For OB, sites within 75 ± 25 µm dorsal to the depth of maximal spike density were selected on each shank, and signals were averaged within and across shanks. Analyses used beta (10–30 Hz), low-gamma (30– 50 Hz), and high-gamma (50–100 Hz) bands.

Spectrograms were calculated using 300 ms Hann windows advanced in 30 ms steps, with spectral power multiplied by frequency to reduce the broadband frequency dependence. Time-resolved band power was calculated by applying third-order Butterworth bandpass filters forward and backward and taking the squared magnitude of the Hilbert analytic signal. Time-resolved OB–PCx magnitude-squared coherence was calculated using 256 ms Hann windows advanced in 16 ms steps. Cross-spectra and autospectra were averaged across matched trials before calculating coherence, which was then averaged across frequency bins within each band.

For scalar comparisons, power and coherence were calculated over a fixed 475 ms interval following baseline or first odor inhalation. This duration corresponded to the measured mean first odor-breath cycle and provided a common analysis interval across recordings. Mean-subtracted LFP segments were analyzed using discrete prolate spheroidal sequence multitaper estimation with a time–bandwidth product of 3 and five tapers. Power spectra were averaged across trials and tapers, multiplied by frequency, and averaged across frequency bins within each band. For coherence, cross-spectra and autospectra were pooled across matched trials and tapers before calculating magnitude-squared coherence and averaging within each band. Each recording–odor pair contributed one scalar value per band and condition. Group comparisons used two-sided Wilcoxon rank-sum tests without correction for multiple comparisons.

### Single unit isolation and analysis

Spike sorting and unit curation were performed using SpikeInterface^55^ and Kilosort4^56^ packages in Python. In brief, raw 30 kHz data were high-pass filtered and bad channels were interpolated prior to spike sorting. Rigid drift correction was applied during spike sorting using Kilosort4’s algorithm (universal template threshold set to 9 and learned template threshold set to 8). In recordings where both dorsal and ventral OB were collected, we split the two regions during spike sorting as they often demonstrated differential drift rates. To curate well-isolated units, we first applied quality control metrics from SpikeInterface’s pipeline ^57,58^ combined with floor cutoffs for unit presence and overall firing rate. We combined this with depth localization for PCx recordings, using a maximal cutoff distance dorsal to the estimated center of the LFP polarity reversal when keeping units for analysis. A unit therefore needed to meet the following criteria for inclusion:

- Inter-spike-interval (ISI) violation ratio < 0.5 (SpikeInterface metric)
- Presence ratio > 0.9 (SpikeInterface metric)
- Amplitude cutoff < 0.1 (SpikeInterface metric)
- Overall firing rate > 0.2 Hz
- Unit location < 350 µm dorsal to the LFP center (PCx only)

Units meeting these criteria were visually inspected to avoid global biases towards under or over-merging units during sorting.

### Optogenetic tagging of PV cells

For opto-tagging recordings, 500 ms, 450 nm light pulses at approximately 10 mW power were delivered at an interval of 4 seconds after the end of odor stimulation to allow for cell identification during post-hoc analysis. Pulse times were aligned and two criteria were applied before calling a unit positive for opto-tagging: 1) firing rate in the pulse window had to exceed firing rate in an equivalent length pre-pulse baseline (threshold of p < 0.001, Wilcoxon rank-sum test) and 2) the unit’s first-spike latency had to be significantly shorter than expected by a homogeneous Poisson null whose rate was estimated from that unit’s own pre-pulse firing rate (binomial test, p < 0.001) and fall within 15 ms of pulse onset. Raster and PSTH plots surrounding pulse windows were also visually inspected to confirm that tagged units responded robustly to light pulses.

### Cell type clustering

Recordings that included optogenetically tagged PV cells (n = 9 recordings, n = 37 cells), all young/old PCx sessions in this manuscript (n = 15) and additional PCx recordings from C57BL/6J mice processed through the same pipeline (n = 37), were pooled and template metrics were calculated for each unit using a combination of custom scripts and built-in SpikeInterface tools:

- Mean ISI
- Burst index (sum of ISIs < 6 ms / sum of ISIs 50-100 ms)
- Main peak to trough ratio
- Pre peak to trough ratio
- Peak to trough duration
- Recovery slope
- Repolarization slope
- Waveform asymmetry

Using this set of features, a grid search of parameters was performed to first project data onto two dimensions using t-SNE and then cluster using DBSCAN. The cluster with the most opto-tagged PV cells was designated as PC_INH._ Additionally, cross-correlograms (CCG) were constructed using SpikeInterface and putative excitatory CCG pairs were identified. Clustering parameters that yielded greater than 20 excitatory pairs in the PC_INH_ cluster were then discarded and outputs were sorted by silhouette score with the top scoring parameters used for clustering in the manuscript. For further validation that the PC_INH_ cluster was predominantly interneurons, we confirmed that the majority of significant inhibitory CCG pairs were co-clustered with the opto-tagged PV cells.

### Inhalation alignment and response measures

A median (15 ms window), Savitzky-Golay (6^th^ order, 125 ms window), and local detrend filter (2 s window, 1.5 s steps) were sequentially applied to the respiration trace and threshold crossings were then used on RMS normalized signal to define the start of inhalations. Unless otherwise stated, all downstream analysis was then aligned with the start of inhalation on the first breath after the start of odor delivery. For first breath response vectors, unit spikes were summed and rate was calculated from the start of the first inhalation of the odor to the start of the second inhalation. For responsivity measures, first breath response vectors were compared to rates during a pre-odor baseline breath, with one-sided Wilcoxon rank-sum tests used to assess whether a given unit was activated or suppressed for an odor stimulus. Percent values were then either averaged across all stimuli by animal or after binning by how many stimuli activated or suppressed a cell before averaging. For PSTH plots, we created a smoothed kernel density function using a Gaussian kernel of 10 ms prior to averaging. For initial peak response magnitudes, we searched for the average index of the peak PSTH value across all animals within the first 75 ms within each region, covering initial rises in population response in both OB and PCx. We then averaged trial values within each population-odor pair ± 2 ms around the peak index value.

### Vector correlations and odor identity decoding

For population vector analysis, first breath responses from all cells were pooled by recording area and responses were trial averaged within stimulus. Trial-averaged response vectors were then compared using the Pearson correlation coefficient. For within animal Pearson’s correlations, response vectors were compiled within an animal, then all trial-to-trial correlations were calculated. For a given odor, the average across stimulus correlation value was subtracted from the average within stimulus value for the final Δ correlation value for each population-odor pair. For within animal odor decoding, response vectors were similarly constructed and split using stratified K-fold (k = 4) cross-validation to test model accuracy in identifying held out trial identities with a linear SVM implemented using the scikit-learn package in Python. Average fold accuracy for a given odor was then reported as a population-odor pair. For pseudopopulation analysis, 50 iterations of cells were randomly sampled at increasing population sizes from all pooled recordings. Δ correlation was calculated as before on the pseudopopulation response array, with averages and error reported across iterations instead of within animal at each cell number step. For decoding analysis, linear SVMs were again constructed, stratified K-fold (k = 4) accuracy was averaged for each iteration as before, and average across iterations was reported at each cell number step.

### Lifetime and population sparseness

Both lifetime and population sparseness were calculated as in previous work ^11,45^. In brief, both measures assess non-uniformity across the measured distribution, with lifetime sparseness calculated using responses of a given unit to the panel of stimuli and population sparseness using all unit responses of a given population within an odor. Values of 1 indicate perfect peakedness (i.e., perfect tuning to a single stimulus for lifetime sparseness or a concentration of response in a single neuron for population sparseness) while values of 0 indicate a completely uniform response. For all measures, first breath response vectors were first trial-averaged within odor. For lifetime sparseness, individual cell sparseness was first calculated, then either plotted to show the distribution by cell or averaged within animal before reporting. For population sparseness, reported values were calculated within each stimulus for a given animal.

### Mean-adjusted variance

To quantify trial-to-trial response variability while accounting for differences in firing rate magnitude, we calculated a mean-adjusted variance for each cell–odor pair. Firing rates were measured on individual trials during the first respiration cycle after odor delivery, and the variance across trials was divided by the corresponding mean firing rate. The same calculation was performed using firing rates during the pre-odor baseline respiration cycle. Mean-adjusted variance values were then averaged across cells within each population–odor pair for statistical comparison between age groups.

### Classifier and principal component analysis on age-related features

Features calculated in prior analysis were first compiled by averaging within animal for any value reported either by cell or by population-odor pair in prior figures. Only features that were significant at the animal or population-odor pair level in prior figures were used in these analyses. Linear SVMs were then constructed to classify age of the animal (binary labels, young or old) using leave-one-out cross validation so that each animal’s age was predicted by a model. Overall accuracy was reported and feature weights were extracted from each SVM and averaged across folds. Accuracy was compared to similarly calculated accuracy on a shuffled distribution of all possible age label recombinations (C(15,7) = 6,435). PCA was then applied to the aging features and feature values were projected onto the first three PCs and the percent of variance explained by each PC was reported. For PCA loading plots, PCA was repeated using leave-one-out splits as with the SVM and average loadings for the top three PCs were reported.

### Statistical tests

Unless otherwise stated with reported p values, two-sided Wilcoxon rank-sum tests were primarily used to assess statistical significance between groups. Welch’s t-test was used to compare recording yields between young and old animals by area. Kolmogorov-Smirnov tests were used to assess differences in the distribution of larger samples of data by individual cells (firing rate and lifetime sparseness tests).

## Data and code availability

All data and code will be made publicly available upon publication.

## Author contributions

I.F.C. and K.A.B. conceived the study. I.F.C. performed experiments and analyzed data. K.A.B. supervised the project. I.F.C. and K.A.B. wrote the manuscript.

## Acknowledgements

We thank Gulce Dikecligil, Joseph Zak, and Valentina Parma for their thoughtful reading of and helpful feedback on the manuscript, and Moona Kim and Wisdom Iwueze for technical support.

