## Supplementary Information for "Aging reshapes odor-evoked dynamics in central olfactory circuits"

| Name | CID | CAS | Diluent | Concentration (% v./v.) |
|---|---|---|---|---|
| Benzaldehyde | 240 | 100-52-7 | Caprylic capric triglyceride | 0.5 |
| Ethyl butyrate | 7762 | 105-54-4 | Caprylic capric triglyceride | 0.5 |
| 1-Hexanol | 8103 | 111-27-3 | Caprylic capric triglyceride | 0.5 |
| 2-Heptanone | 8051 | 110-43-0 | Caprylic capric triglyceride | 0.5 |
| Heptaldehyde | 8130 | 111-71-7 | Caprylic capric triglyceride | 0.5 |
| Acetophenone | 7410 | 98-86-2 | Caprylic capric triglyceride | 0.5 |
| 2-pentanone | 7895 | 107-87-9 | Caprylic capric triglyceride | 0.5 |
| Butyric acid | 264 | 107-92-6 | Caprylic capric triglyceride | 0.5 |
| Isoamyl acetate | 31276 | 123-92-2 | Caprylic capric triglyceride | 0.5 |
| 6-methyl-5-hepten-2-one | 9862 | 110-93-0 | Caprylic capric triglyceride | 0.5 |
| Isobutyl acetate | 8038 | 110-19-0 | Caprylic capric triglyceride | 0.5 |
| Methyl hexanoate | 7824 | 106-70-7 | Caprylic capric triglyceride | 0.5 |
| Nerol | 643820 | 106-25-2 | Caprylic capric triglyceride | 0.5 |
| 2-Butanone | 6569 | 78-93-3 | Caprylic capric triglyceride | 0.5 |
| R-(+)-Limonene | 440917 | 5989-27-5 | Caprylic capric triglyceride | 0.5 |
| Caprylic capric triglyceride | 90474509 | 73398-61-5 | Caprylic capric triglyceride | 100 |

### Supplementary Table 1 – Stimulus panel

| Session ID | OB Units | PC units | $PC_{EXC}$ | $PC_{INH}$ | Recording Age (days) | Recording Age (months) |
|---|---|---|---|---|---|---|
| 040281__25_08_21 | 87 | 556 | 505 | 51 | 108 | 3.55 |
| 040282__25_08_20 | 27 | 357 | 335 | 22 | 107 | 3.52 |
| 040283__25_09_03 | 118 | 283 | 261 | 22 | 121 | 3.98 |
| 040294__25_09_24 | 102 | 472 | 449 | 23 | 103 | 3.39 |
| 040343__25_12_09 | 75 | 387 | 360 | 27 | 106 | 3.48 |
| 040344__25_12_12 | 40 | 184 | 173 | 11 | 109 | 3.58 |
| 040345__25_12_16 | 85 | 312 | 287 | 25 | 113 | 3.72 |
| 040088__25_12_24 | 59 | NaN | NaN | NaN | 581 | 19.1 |
| 040089__25_12_18 | 55 | 277 | 259 | 18 | 575 | 18.9 |
| 9F10T-2_BEC__25_08_08 | 111 | 566 | 506 | 60 | 612 | 20.12 |
| 9F10T-2_NEC__25_08_12 | 107 | 406 | 359 | 47 | 616 | 20.25 |
| 9F8F_2_LEC__25_11_23 | 62 | 241 | 215 | 26 | 770 | 25.32 |
| 9F8F_2_NEC__25_12_05 | 36 | 185 | 180 | 5 | 782 | 25.71 |
| 9F8F_2_REC__25_11_22 | 24 | 277 | 260 | 17 | 769 | 25.28 |
| 9F9M_LEC__25_09_04 | 103 | 213 | 200 | 13 | 686 | 22.55 |
| 9F9M_NEC__25_08_29 | 67 | 237 | 203 | 34 | 680 | 22.36 |

### Supplementary Table 2 – Animal ages and recording yields

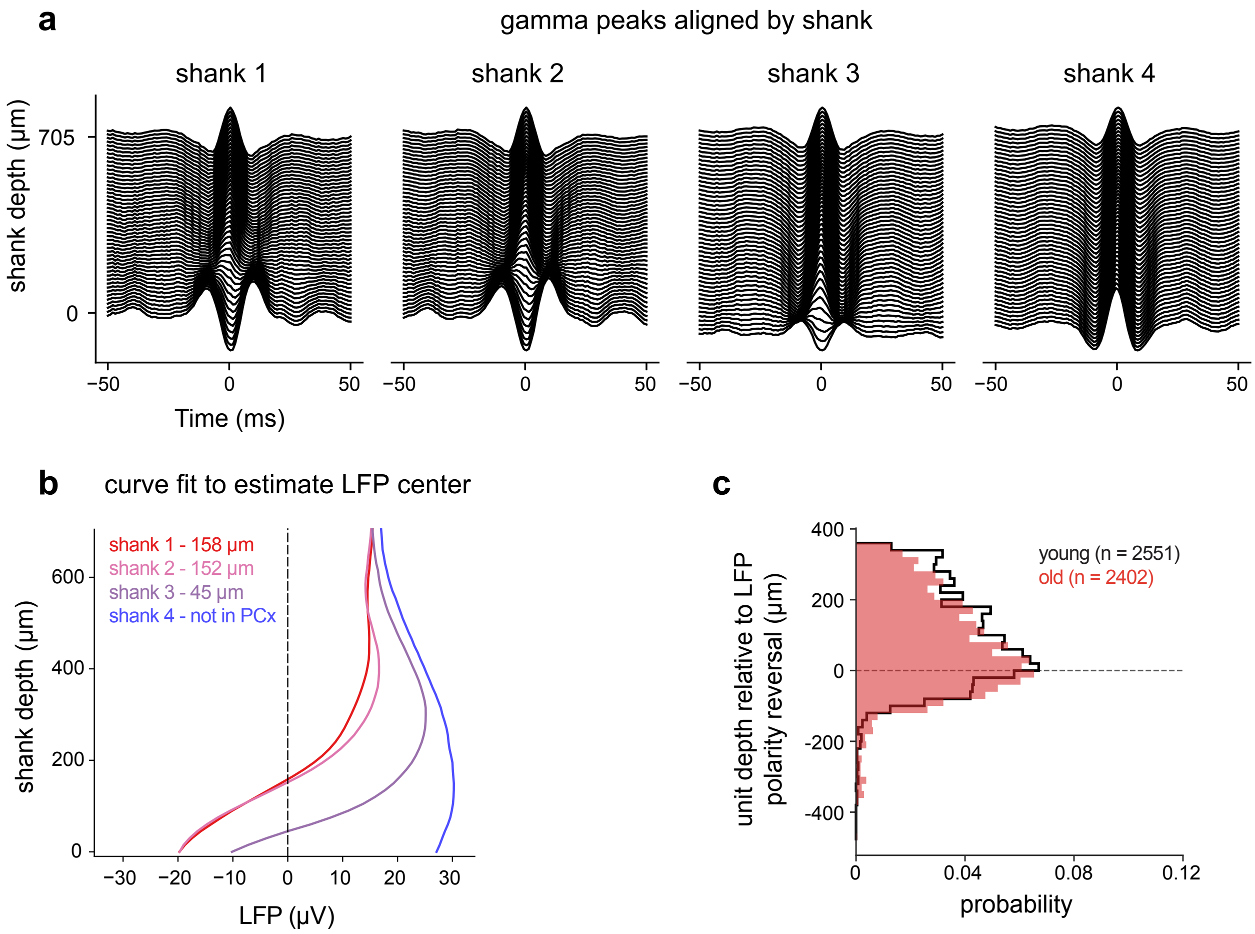


**Supplementary Figure 1. PCx probe localization and distribution of recorded units across depth**

**a** Gamma-band event-triggered average LFP traces from an example PCx probe, aligned to detected high-amplitude gamma-band events and ordered by channel depth on each shank**. b** LFP amplitude at the gamma-event peak plotted as a function of depth for each shank. Estimated polarity-reversal depths are shown for shanks 1–3, consistent with the probe spanning PCx; shank 4 lacked a detectable polarity reversal and was excluded from analysis. **c** Probability-normalized distributions of PCx unit depths relative to the estimated LFP polarity reversal for young and old mice.

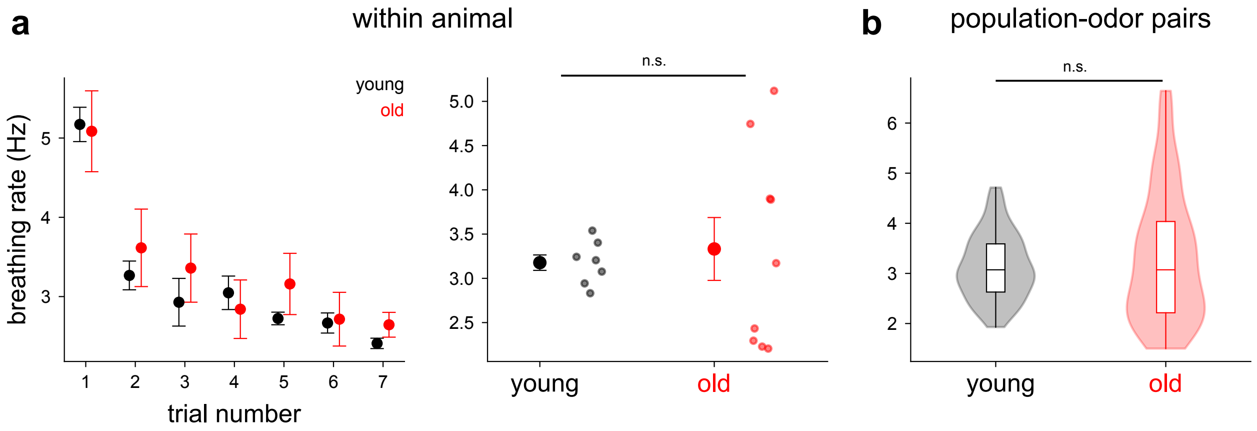


**Supplementary Figure 2. Respiration rate during odor presentations**

**a** Breathing rate during the 2 s following odor delivery, shown across trial number (left) and with all trials pooled (right). Mean ± SEM across animals. Points are individual animals. **b** Distribution of breathing rates across population-odor pairs, calculated over the same 2-s period. Violin plots show the distribution of values, with embedded box plots indicating the median and interquartile range. Population-odor pair and animal-level comparisons were performed using Wilcoxon rank-sum tests; n.s., not significant.

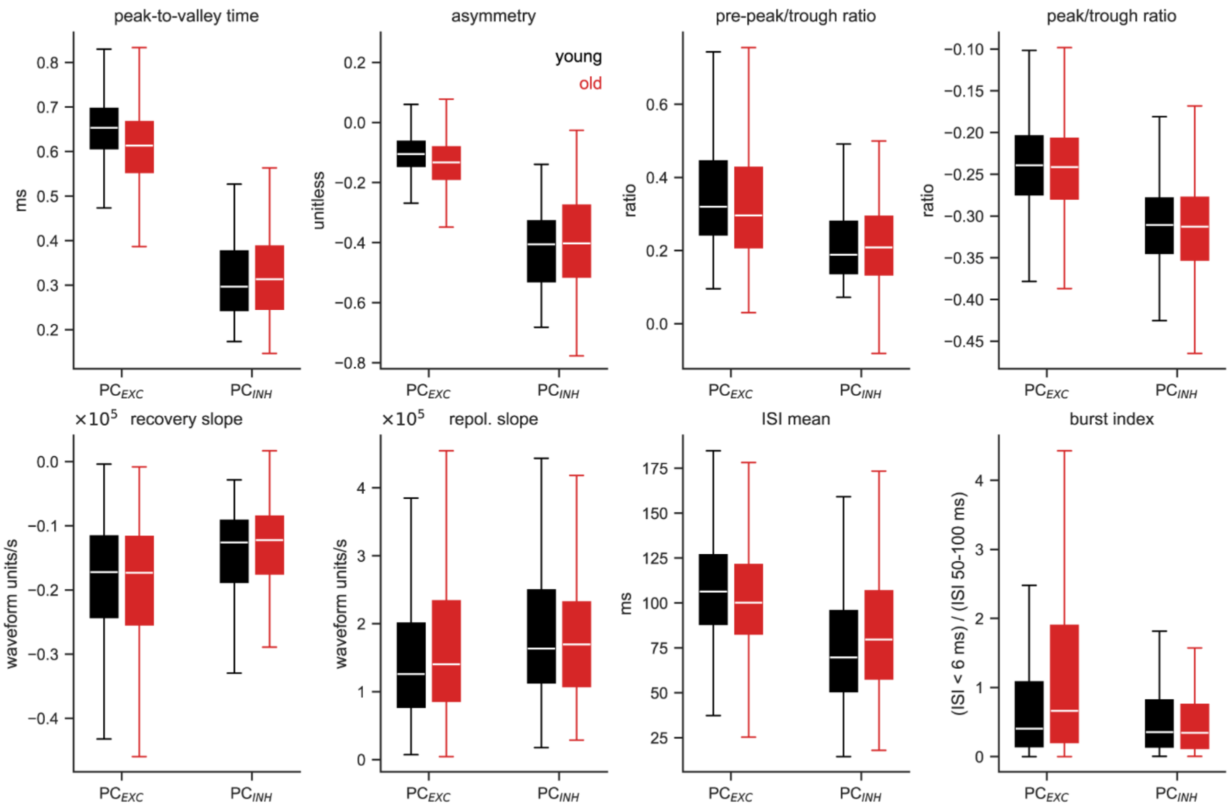


**Supplementary Figure 3. Waveform and spiking features used for PCx cell-type clustering**

Distributions of waveform and spiking features for putative $PC_{EXC}$ and $PC_{INH}$ cells in young (black) and old (red) mice. Features shown are peak-to-valley time, waveform asymmetry, pre-peak/trough ratio, peak/trough ratio, recovery slope, repolarization slope, mean inter-spike interval (ISI), and burst index, defined as the ratio of ISIs < 6 ms to ISIs between 50–100 ms. Box plots indicate the median and interquartile range; whiskers show the data range.

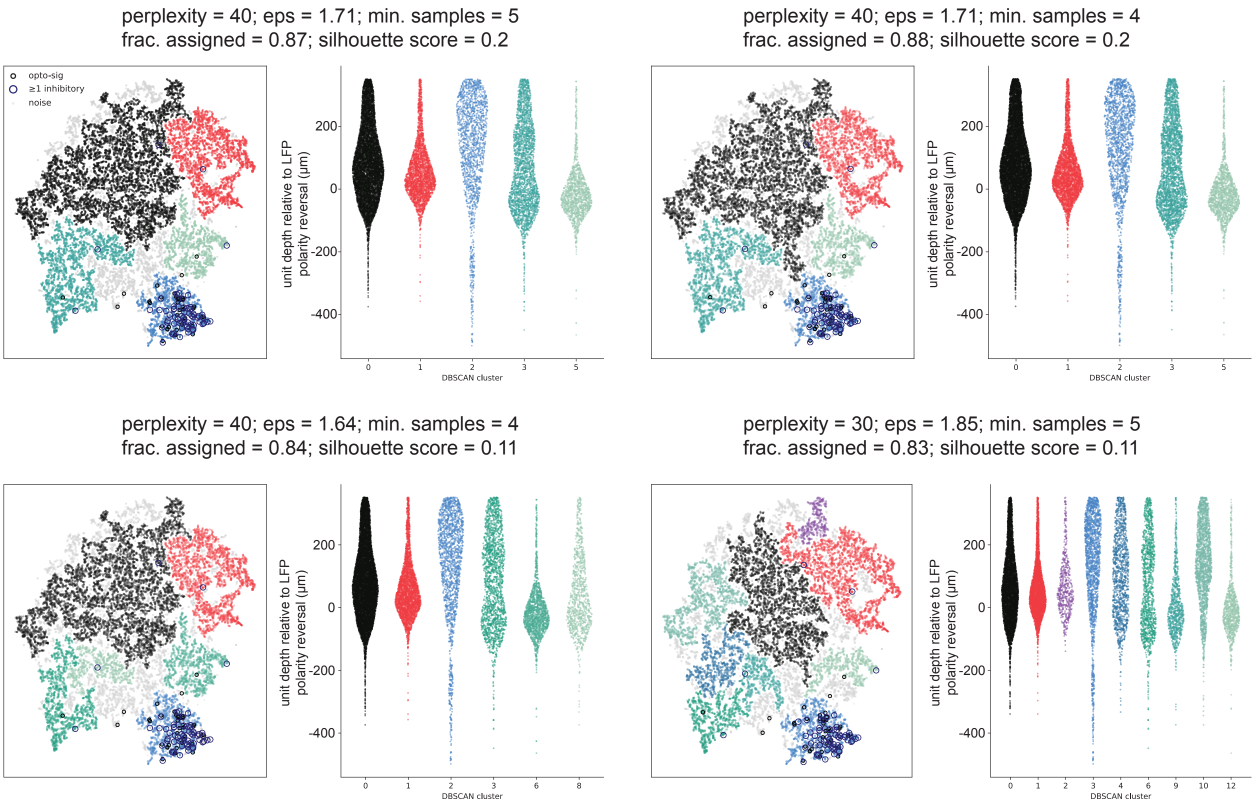


**Supplementary Figure 4. Robust identification of the $PC_{INH}$ cluster across clustering parameters**

Example t-SNE projections and DBSCAN clustering solutions from the four highest-scoring parameter combinations, ranked by silhouette score after a parameter sweep across a small set of t-SNE and DBSCAN settings. Optogenetically tagged PV cells and units with putative inhibitory CCG interactions are indicated on each projection. Adjacent plots show the distribution of unit depths relative to the PCx LFP polarity reversal for each DBSCAN cluster. Across these top-scoring parameter combinations, the number and organization of non-inhibitory clusters varied, whereas opto-tagged PV cells and putative inhibitory units consistently occupied a distinct cluster with a characteristic depth distribution, supporting robust identification of the $PC_{INH}$ population.

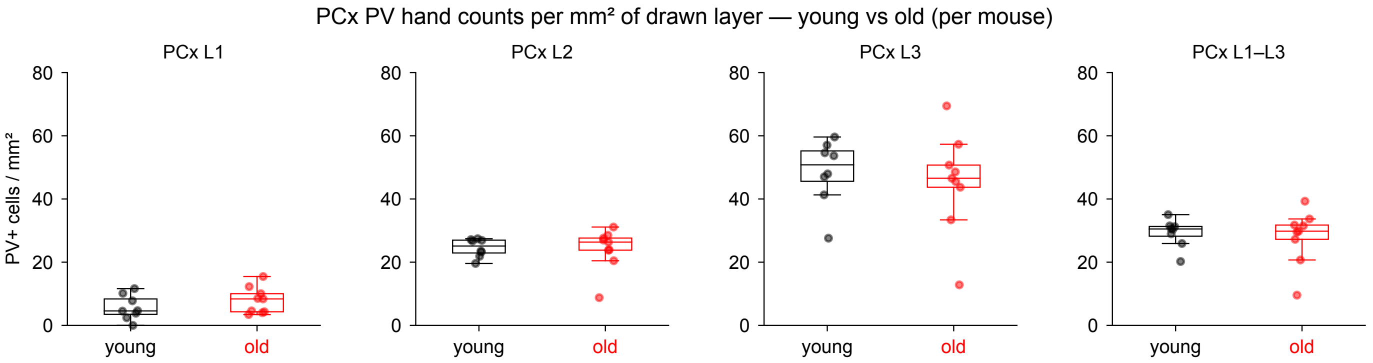


**Supplementary Figure 5. Parvalbumin-positive cell density in anterior piriform cortex is unchanged with age.**

Density of parvalbumin-positive (PV+) cells in layers 1, 2, and 3 of anterior piriform cortex (PCx), and across all three layers combined, in young (black) and old (red) mice. Each point represents the PV+ cell density (cells/mm²) for one animal. PV+ cell density was quantified in all animals from the aging cohort for which histology was available ($n = 8$ young, 9 old), including one young animal that was recorded with a different odor panel and therefore not included in the electrophysiological analyses. PV+ cell density did not differ significantly between age groups in layer 1 ($p = 0.370$), layer 2 ($p = 0.541$), layer 3 ($p = 0.606$), or across all layers ($p = 0.888$; two-sided Wilcoxon rank-sum tests). Boxplots indicate the median and interquartile range, with whiskers extending to 1.5 times the interquartile range.
